## Supplemental figures for "Persistence of intact HIV-1 proviruses in the brain during antiretroviral therapy"

Figure S1

A

Clinical and demographical data of study participants

| Participant | Gender | Age of death | Duration of HIV-1 infection (yr) from diagnosis date | Time on HAART (yr) | HAART regimen | CD4 count before death (cells/ul) | Viral load before death (copies/ml) |
|---|---|---|---|---|---|---|---|
| 1 | Female | 38 | 16 | 16 | FTC, RPV, TAF | 1625 | Undetectable |
| 2 | Male | 68 | 25 | 1 | FTC, TAF, BIC | 165 | Undetectable |
| 3 | Male | 52 | 1 | 1 | BIC, FTC, TAF | 145 | 136 |

B

Cell numbers analyzed from each tissue of each study participant

| | Compartment | Participant 1 (million cells) | Participant 2 (million cells) | Participant 3 (million cells) |
|---|---|---|---|---|
| Brain tissues | Basal ganglia | 67.25 | 18.06 | 32.85 |
| | Thalamus | 35.81 | 5.77 | 21.83 |
| | Occipital lobe | 87.17 | 29.07 | 43.65 |
| | Frontal lobe | 64.29 | 13.39 | 64.91 |
| | Periventricular white matter | | | 36.42 |
| Non-brain tissues | Lymph nodes | 28.86 | 15.57 | |
| | Spleen | 63.69 | 104.15 | |
| | Colon | 54.02 | 17.74 | |
| | Liver | 72.91 | 43.59 | |
| | Pancreas | 91.36 | 65.1 | |
| | Terminal ileum | | 1.46 | |
| | Kidney | 57.02 | 25.9 | |
| | Ovary | 51.96 | | |
| | Uterus | 79.11 | | |
| | Testes | | 4.74 | |
| | Prostate | | 19.5 | |
| | Thyroid | 53.65 | 46.63 | |
| | Adrenal | 39.43 | 14.87 | |
| | Total | 846.53 | 425.54 | 199.66 |

Figure S2

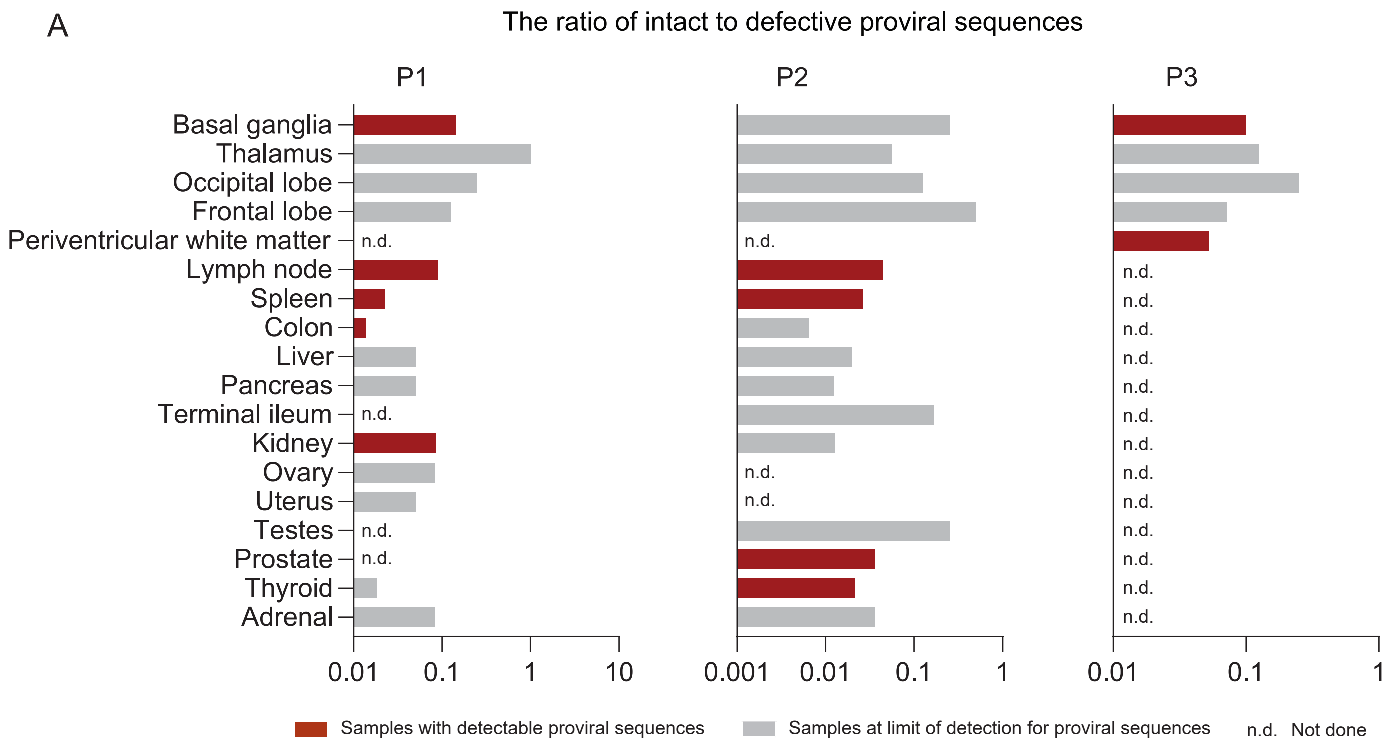
A
The ratio of intact to defective proviral sequences
P1
P2
P3
Basal ganglia
Thalamus
Occipital lobe
Frontal lobe
Periventricular white matter
Lymph node
Spleen
Colon
Liver
Pancreas
Terminal ileum
Kidney
Ovary
Uterus
Testes
Prostate
Thyroid
Adrenal
n.d.
0.01 0.1 1 10
0.001 0.01 0.1 1
0.01 0.1 1
Samples with detectable proviral sequences
Samples at limit of detection for proviral sequences
n.d. Not done

Figure S3

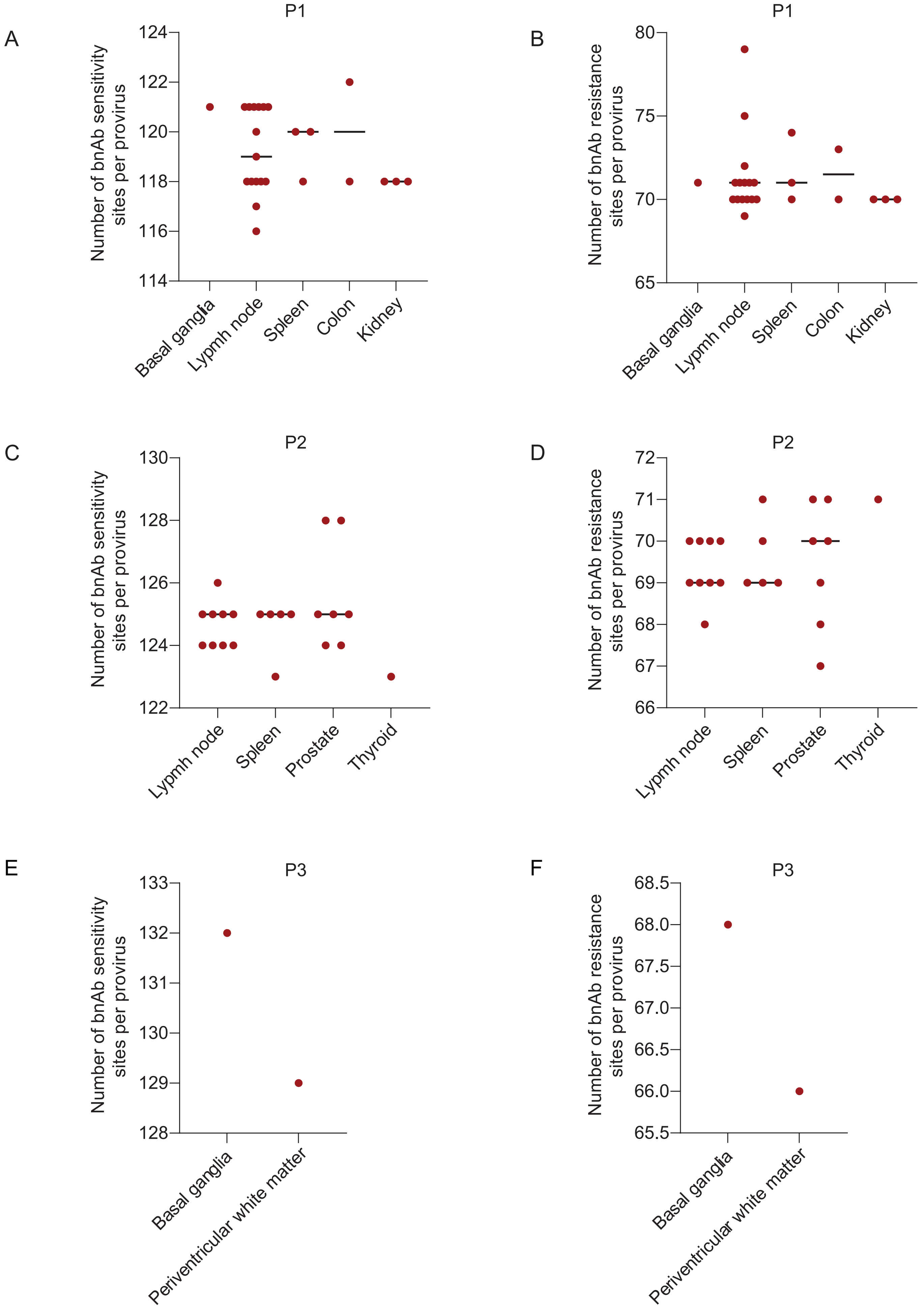
A
P1
Number of bnAb sensitivity sites per provirus
124
122
120
118
116
114
Basal ganglia
Lypmh node
Spleen
Colon
Kidney
B
P1
Number of bnAb resistance sites per provirus
80
75
70
65
Basal ganglia
Lypmh node
Spleen
Colon
Kidney
C
P2
Number of bnAb sensitivity sites per provirus
130
128
126
124
122
Lypmh node
Spleen
Prostate
Thyroid
D
P2
Number of bnAb resistance sites per provirus
72
71
70
69
68
67
66
Lypmh node
Spleen
Prostate
Thyroid
E
P3
Number of bnAb sensitivity sites per provirus
133
132
131
130
129
128
Basal ganglia
Periventricular white matter
F
P3
Number of bnAb resistance sites per provirus
68.5
68.0
67.5
67.0
66.5
66.0
65.5
Basal ganglia
Periventricular white matter
